# Oncogenic STIL-mediated loss of BRCA1 functionality causes DNA damage and centrosome amplification

**DOI:** 10.1101/2024.02.11.579865

**Authors:** Srishti Sanghi, Priyanka Singh

**Affiliations:** Department of Bioscience & Bioengineering, Indian Institute of Technology Jodhpur NH 62, Nagaur Road, Karwar 342037, Jodhpur, Rajasthan, India

**Keywords:** STIL, BRCA1, Centrosome, Breast Cancer, Cell division

## Abstract

DNA damages increase centrosome number, a typical phenotype in many cancers. However, the molecular linkers between the two pathways remain elusive. STIL (SCL/TAL1 Interrupting locus) is a core centrosome-organizing protein overexpressed in different cancer types. This work shows that STIL forms a complex with the DNA damage response protein, BRCA1, and is required for regulating its protein stability. Subsequently, STIL depletion resulting in BRCA1 loss from the nucleus causes centrosome amplification and DNA damage. We also identified a heterozygous missense mutation (S76L) in STIL from the pan-cancer datasets, which does not affect total BRCA1 protein levels. However, it reduces BRCA1 nuclear localization with enhanced centrosome recruitment simultaneously at the S-phase. The enhanced BRCA1 at centrosome causes an increase in Aurora-A kinase levels at centrosomes, thus culminating in DNA damage and centrosome amplification. The expression of exogenous BRCA1 can push back this imbalance of the STIL-BRCA1 axis in oncogenic conditions. The amplified centrosomes in oncogenic STIL condition cluster in the M phase and cause pseudo-bipolar spindle organization. We could decluster them by the HSET(KIFC1)-specific inhibitor, CW069, suggesting a promising target for anti-cancer treatment.

## INTRODUCTION

The gene for STIL (SCL/TAL1 Interrupting locus) was first cloned from the chromosomal rearrangements in T-cell leukemia^1^. STIL is required for cell growth and proliferation, as its loss of function mutations are lethal in mice at midgestation^2,3^ and zebrafish^4^. Enhanced expression of STIL is observed in different cancers^5,6^, and it has been established as an oncogene^7^.

STIL is mainly a cytoplasmic protein, which is localized at centrosomes. However, a small percentage of protein is also detected in the nucleus^8^. A centrosome contains an orthogonally arranged pair of microtubule-based cylindrical structures at its core called the centrioles, which are embedded in a protein-rich matrix referred to as the pericentriolar material (PCM). Centrosome duplicates during the S-phase of the cell cycle, which involves a cartwheel-like arrangement of core centriole proteins (procentriole) adjacent to each preexisting centriole. STIL is one of the core centriole duplication factors, whose functional orthologues are SAS5 in worms^9^ and ANA2 in flies^10^. Human STIL contains several conserved regions^11^ (**Figure S1A**), which include a Proline-rich *<u>C</u>*onserved *<u>R</u>*egion (CR2, 385-499 residues) present at the N-terminal region and involved in interactions with another core centriole protein, CPAP/CENPJ^12^. An extended CR1 region (26-324 residues) is explicitly found in vertebrates, but little is understood about its involvement in STIL functioning. The middle coiled-coil region (721-748 residues) interacts with the centrosome kinase, PLK4^13^, or the mitotic kinase CDK1, to regulate their centriole localization^14^. A patch of hydrophobic residues in the coiled-coil region is involved in oligomerization, which is required for *de novo* centriole organization^15^. The C-terminal region of STIL has a STAN domain (1052-1148 residues), which interacts with the centriole cartwheel spoke protein SAS6^13^ and helps in procentriole organization (**Figure S1A**). The C-terminal KEN box in STIL is required for promoting APC/C (Anaphase Promoting Complex/Cyclosome)-mediated protein degradation at mitotic exit^16^. Consequently, STIL depletion inhibits centriole duplication^12^, and its overexpression triggers supernumerary centrioles^17^.

In the 19^th^ century, Theodor Boveri (1862-1915) postulated the involvement of extra centrosomes in developing malignant tumors^18^. Thereafter, an increase in centrosome number is found to be positively correlated with aneuploidy and chromosomal instability in different tumor types. Extra centrosomes in *Drosophila* can affect asymmetric divisions of larval neural stem cells, and transplantation of such aberrant cells in the abdomen of host flies induces tumor formation^19^. Besides, amplified centrosomes are observed in many precancerous and preinvasive lesions in humans, suggesting their involvement during the early stages of tumor progression^20–24^. Due to the ubiquitous presence of supernumerary centrosomes in different tumor types, it is now considered one of the “hallmarks” of cancer with a potential prognostic value^25,26^.

DNA damages activate nuclear ATM/ATR-dependent damage response signaling pathways. Outside the nucleus, these damage response proteins are reported at centrosomes, making it a platform for crosstalk between nuclear-cytoplasmic signaling. The breast and ovarian cancer susceptibility gene, BRCA1 (Breast cancer gene 1), is a tumor suppressor that binds to DNA. During the S-phase, BRCA1 displays nuclear puncta, representing different nuclear complexes that protect stalled DNA replication forks from degradation^27^, silencing of inactive X-chromosome^28^, chromatin modifications, and epigenetic silencing^29^ under various conditions. BRCA1 contains a RING domain at the N-terminal and two tandem BRCT domains at the C-terminus region. BRCA1 and BARD1 (BRCA1-associated RING domain protein 1) heterodimerizes through their respective RING domains. The stable complex of BRCA1-BARD1 interacts with other accessory proteins for DNA damage repair^30^ by homologous recombination^31^ and centrosome regulation^32–35^. BRCA1 can shuttle between the nucleus and cytoplasm to carry such diverse cellular activities.

The BRCT regions of BRCA1 are also required for its localization at centrosomes^36^, where it interacts with γ-tubulin. BRCA1-dependent ubiquitination of γ-tubulin inhibits centrosome aster formation at the S-phase^37^. However, in the M-phase of the cell cycle, Aurora-A/B kinase-mediated phosphorylation of BRCA1 inhibits its ubiquitination activity^38^, thus allowing microtubule nucleation. BRCA1 mutations that cause loss of functionality exhibit centrosome amplification; however, it is still unclear whether the BRCA1-mediated ubiquitin ligase or nuclear signaling regulates centrosome duplication in such cases. The BRCT domain of BRCA1 recognizes and binds to its interactors *via* a unique S-X-X-F motif, where Serine (S) is phosphorylated^39^. This motif has been identified in DNA double-strand repair proteins, Abraxas-Rap80^40^, BRIP1 (BACH1/FANC)^41^, and CtIP^42^. BRCA1 can also interact with the core centrosome protein STIL^43^, but little is understood about the interplay between STIL and BRCA1.

This work provides evidence that STIL and BRCA1 crosstalk with each other to maintain DNA integrity and centrosome numbers. We further show the relevance of this STIL-BRCA1 axis in cancer.

## RESULTS

### Double-stranded DNA damages cause centrosome amplification

Centrosomes provide a platform for DNA damage response (DDR) signaling, and many DDR proteins, including BRCA1, also localize at centrosomes^36^. We also show that double-stranded DNA damage inflicted by UV treatment of MCF7 cells causes a significant (55.5%) increase in the population of S-phase cells with amplified centrosome numbers. This phenotype was rescued on STIL depletion (**Figure 1A,C**), suggesting that STIL is an intriguing linker between DNA damage and centrosome amplification. The DDR protein, BRCA1 depletion, or overexpression also causes 20-30% of cells with amplified centrosomes (>4 centin-1 foci per cell) at the S-phase (**Figure 1B,D**). STIL interacts with BRCA1^43^ and also serves as a core centrosome organizing protein. Accordingly, we also show, by immunoprecipitation, that endogenous STIL and BRCA1 are part of the same complex in S-phase MCF7 cells (**Figure 1E**). Although BRCA1 is mainly a nuclear protein, it co-localizes with STIL at centrosomes during the interphase (**Figure S1B**).

**Figure 1:**
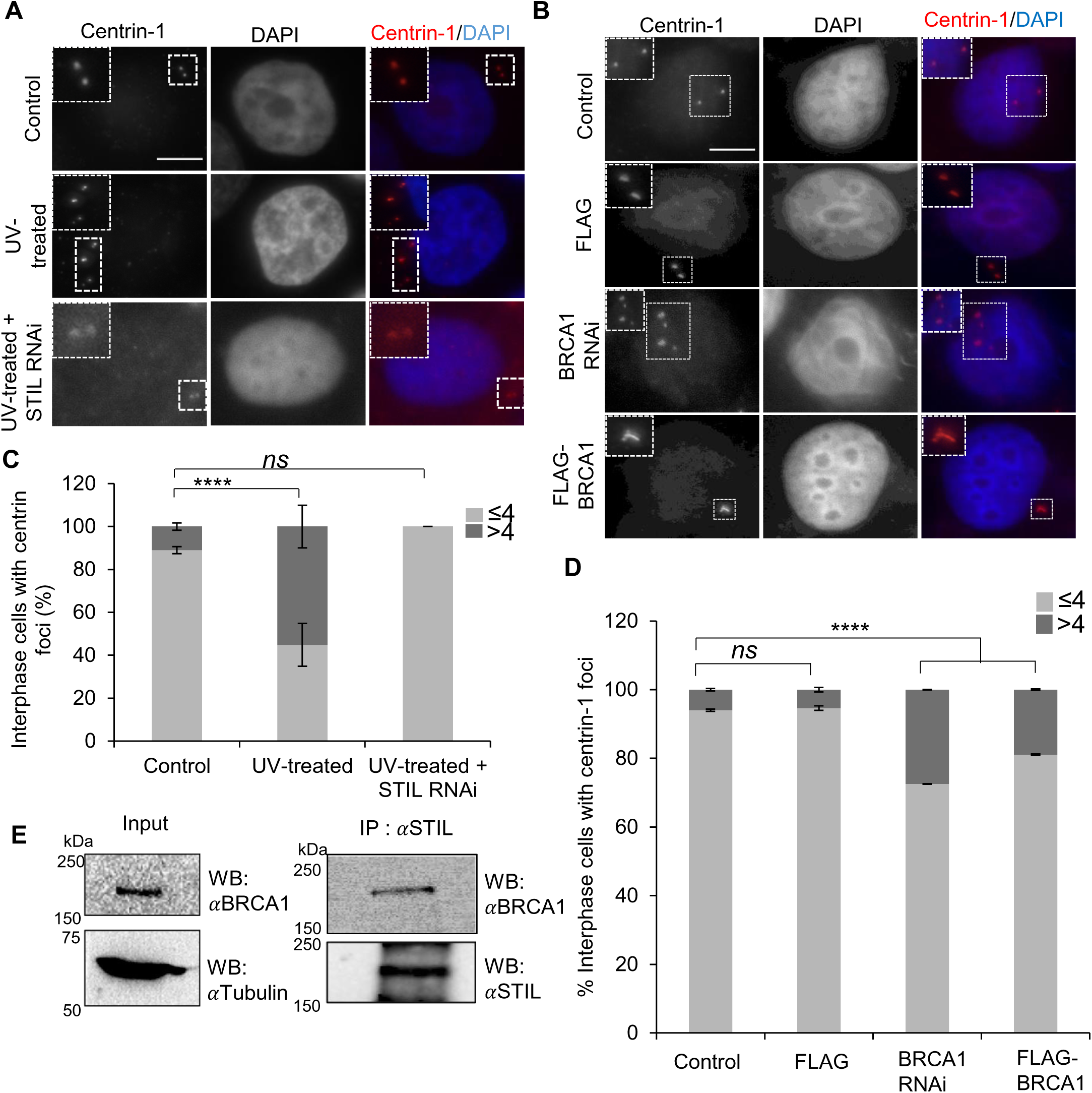
DNA damages affect centrosome number. (**A**) Immunofluorescence images of S-phase synchronized MCF7 cells stained for centrin-1 (centriole marker, red in merge) and DAPI (nuclear marker, blue in merge) in control, UV-treated and UV-treated with STIL depleted using siRNA (STILRNAi) conditions. Insets show a magnified view of centrioles. Scale bar: 5μm. (**B**) Immunofluorescence images for S-phase synchronized MCF7 cells stained for centrin-1 and DAPI. Insets show a magnified view of the centrioles. Scale bar: 5μm. (**C**) Bar graph showing the average percentage of interphase cells±SEM with >4 (dark gray) and ≤4 (light gray) centrin-1 dots for indicated conditions in (A). Results are from two independent experiments (n=80-100). ****p<0.0001; ns, not significant (Chi-squared test). (**D**) Bar graph showing the mean percentage of interphase cells ±SEM with >4 (dark gray) and ≤4 (light gray) centrin-1 dots for indicated conditions in (B). Results are from two independent experiments (n=100–150). ns, not significant (p>0.05); ****p<0.0001 (Chi-squared test). (**E**) Western blot showing immunoprecipitation (IP) of endogenous STIL from MCF7 cells, which was developed (WB) using STIL and BRCA1 antibodies. The blot is a representation of two repeats.

### STIL regulates BRCA1 protein stability to maintain DNA integrity

We further observed that depletion of STIL by siRNA (**Figure S1C**) resulted in a decrease of total BRCA1 protein (**Figure 2A**), suggesting a requirement of STIL for BRCA1 protein stability. Subsequently, we found enhanced ubiquitination of BRCA1 (**Figure 2B**) and a reduced number of BRCA1 nuclear punta at S-phase on STIL depletion (**Figure 2C,D)**. On the other hand, BRCA1 depletion (**Figure S1D**) does not affect the amount of immunoprecipitated STIL protein from the total cell extracts (**Figure 2E**), and STIL could localize at the centrosomes (**Figure 2F**). We found an activation of the p53-p21 pathway on BRCA1 depletion (**Figure 2G**). Accordingly, we show by comet assay that upon STIL depletion where BRCA1 is degraded (**Figure 2A**) and BRCA1 depletion both cause significant DNA damage compared to control cells, as scored by the percentage of DNA in the comet tail (**Figure 2H,I**).

**Figure 2.**
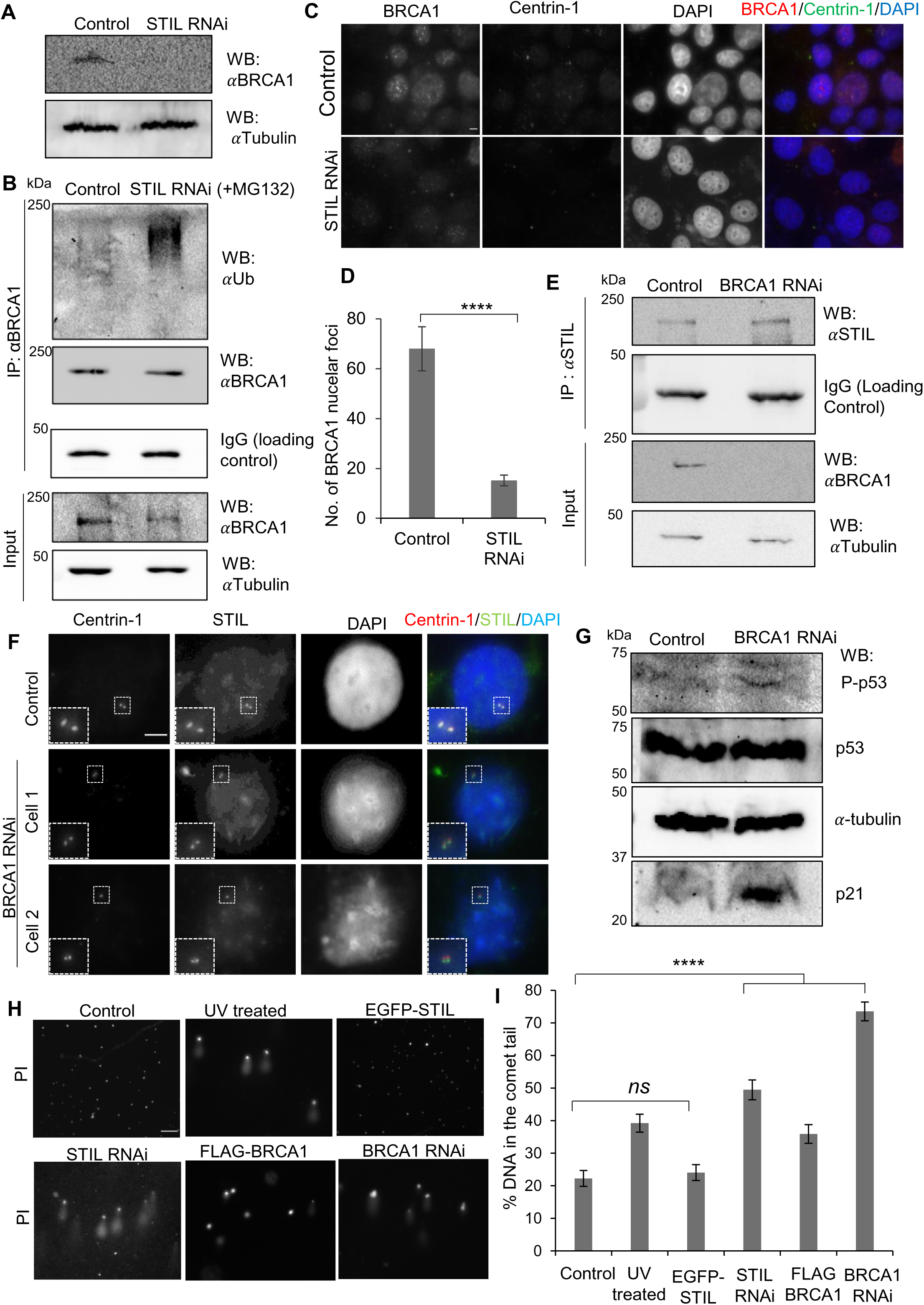
STIL-BRCA1 axis is involved in DNA damage. (**A**) Western blot showing endogenous BRCA1 levels in the total cell lysates of MCF7 cells in control *vs*. STIL RNAi α-tubulin serves as loading control. The blot is representative of four repeats. (**B**) Western blot showing inputs and ubiquitination levels of immunoprecipitated (IP) endogenous BRCA1 in control *vs*. STIL depletion (RNAi) condition. IgG bands serve as loading control for IP and α-tubulin as loading control for input. The blot is representative of four repeats. (**C**) Immunofluorescence images of MCF7 cells stained for endogenous BRCA1 and centrin-1 (centrosome marker). Scale bar: 5μm. (**D**) Bar graph showing the average number of BRCA1 nuclear foci per cell±SEM for control *vs.* STIL RNAi conditions. Results are from two independent experiments (n=70-100). **** p<0.0001 (Two-tailed unpaired Student’s t-test). (**E**) Western blot showing amount of immunoprecipitated (IP) in control vs. BRCA1 depleted (RNAi) conditions. The blot is representative to two repeats. Loading control for IP is IgG and α-tubulin for the inputs. (**F**) Immunofluorescence images for S-phase synchronized MCF7 cells control *vs*. BRCA1 RNAi. Cells are stained for endogenous STIL, centrin-1 (centriole marker), and DAPI (nuclear marker). (**G**) Western blot showing the total amount of phosphorylated p53 (P-p53), p53 and p21 levels in control *vs*. BRCA1 depletion (RNAi) condition. α-tubulin bands serve as loading control. The blot is representative of two repeats. (**H**) Immunofluorescence images of MCF7 cells for comet assay and stained with propidium iodide (PI) to assess varying levels of DNA damage for different conditions. Scale bar: 100μm. (**I**) Bar graph showing the average percentage of DNA in comet tail±SEM as quantified from the experiment (H). Results are from two independent experiments (n=150-250). ns, not significant (p>0.05); ****p<0.0001 (Two-tailed unpaired Student’s t-test).

### Identification of STIL S76L oncogenic mutation in the CR1 region

To understand the significance of the STIL-BRCA1 network in cancer, we analyzed pan-cancer mutations in the *STIL* gene available on public databases, including Cancer Genome Atlas (TCGA) (https://www.cancer.gov/tcga), Catalogue Of Somatic Mutations In Cancer (COSMIC)^44^, and International Cancer Genome Consortium (ICGC)^45^. We found that 45% of cancer-associated mutations in STIL are missense. The CR1 region was found to be frequently mutated, with 141 reported mutations (20%) (**Figure 3A**). The expression of EGFP tagged STIL (EGFP-STIL, 1-1287 residues) in breast cancer cells, MCF7, resulted in only a subtle increase (2.5 folds) in a population of cells with centrosome amplification (>2 γ-tubulin foci per cell) as scored using the pericentriolar marker, γ-tubulin in prophase cells (**Figure S1E,F)**. However, the expression of the N-terminal of STIL (1-619 residues) caused a 7.6-fold increase in cell population with amplified centrosomes, which is even more as compared to the C-terminal region (5.5 folds) of STIL encompassing 619-1287 residues. This corroborates the bioinformatics analysis and suggests an essential role of the STIL N-terminal region in regulating its oncogenicity. Further analysis of CR1-associated cancer mutations led us to narrow down a frequently reported missense mutation, which changes the serine residues at the 76^th^ position to leucine (S76L). The heterozygous STIL-S76L was reported in different epithelial cancer types, including skin, stomach, colon, and endometrial cancer (**Figure 3A,B)**.

**Figure 3:**
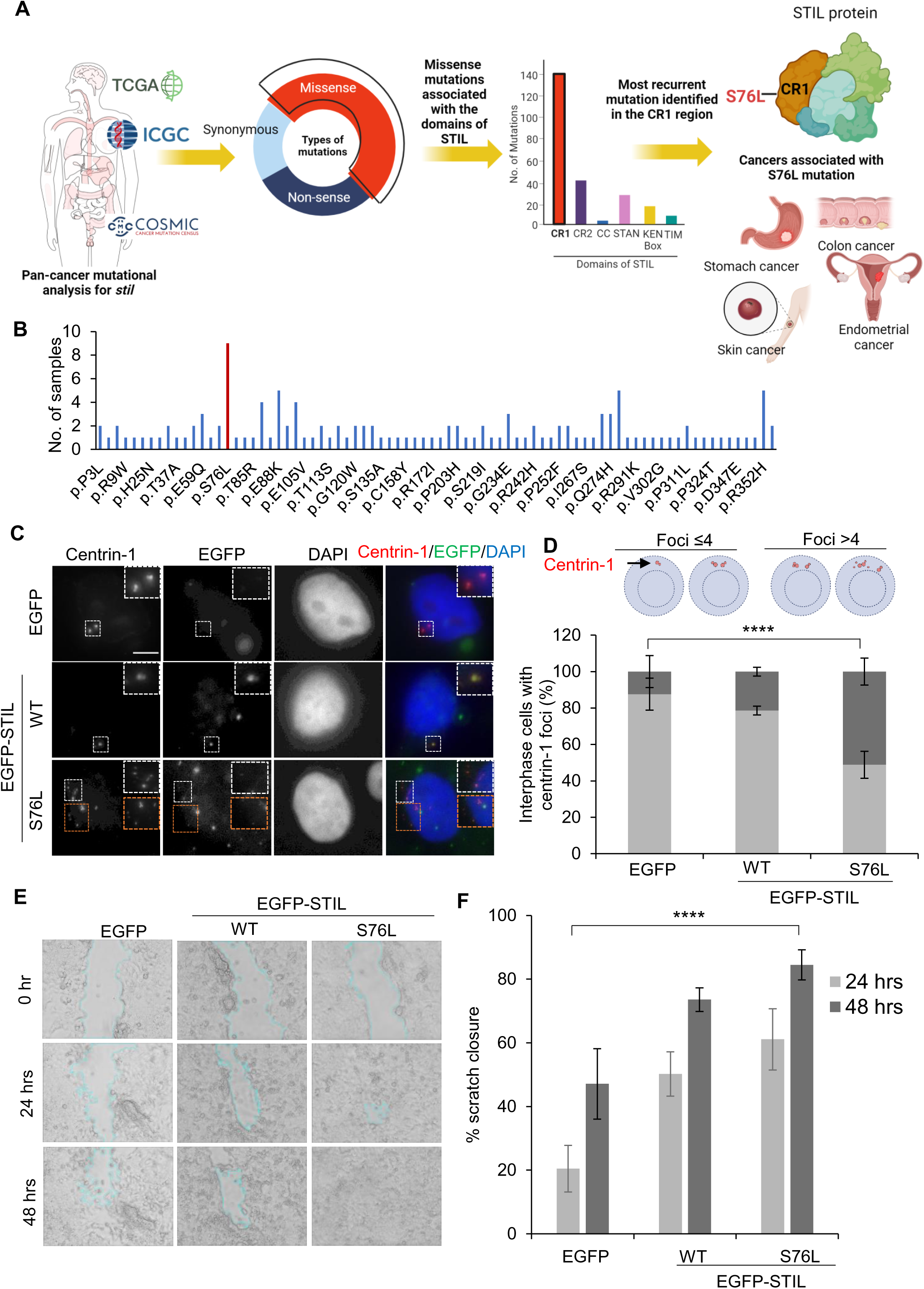
Oncogenic STIL (S76L) is associated with cancer phenotypes. (**A**) The workflow for identifying STIL-associated cancer mutation from the pan-cancer datasets available in various cancer databases. The majority of missense mutations were reported in the CR1 region of STIL. (**B**) The bar graph represents the number of reported missense mutations associated with each residue in the CR1 region of STIL. (**C**) Immunofluorescence images of S-phase synchronized MCF7 cells expressing EGFP (control) or EGFP-STIL (wild type, WT; S76L, mutant). The cells were stained with centrin-1 (centriole marker) and DAPI (nuclear marker). Insets show a magnified view of centrioles. Scale bar: 5μm. (**D**) The top panel shows a cartoon depicting interphase cells scored under ≤4 or >4 centrin-1 foci. Bar graph showing the mean percentage of interphase cells±SEM with >4 (dark gray) and ≤4 (light gray) centrin-1 foci for the respective conditions in (C). Results are from two independent experiments (n=100–180). ****p<0.0001 (Chi-squared test). (**E**) Bright-field images showing scratch closure at 0 hr, 24 hrs, and 48 hrs in MCF7 cells transfected with EGFP (control) or EGFP-STIL (wild type, WT; S76L, mutant). (**F**) Bar graph showing the mean percentage of scratch closure±SEM at 24 hrs (light gray) and 48 hrs (dark gray) for conditions in (E). The results are from two independent experiments. ****p<0.0001 (two-tailed unpaired Student’s t-test).

The expression of EGFP-STIL-S76L in the MCF7 cells resulted in a significant (70.8%) population of prophase cells with amplified centrioles (>4 centrin-1 foci per cell; a distal centriole marker), compared to control cells expressing only EGFP (**Figure S2A,B**). Centriole duplicates at the S-phase of the cell cycle, so S-phase synchronized cells expressing EGFP-STIL-S67L already show a significant population (51%) with amplified centrosomes (>4 centrin-1 foci per cell) (**Figure 3C,D**). That suggests that the S76L mutation affects the centriole duplication function of STIL protein. Next, we performed a scratch assay to analyze cell migration, which is linked with tumorigenesis^46^. The MCF7 cells expressing EGFP-STIL-S76L closed the scratch by 84.5% in 48 hrs, whereas in control (EGFP expressing cells), only 47% scratch closure was observed (**Figure 3E,F)**, indicating that the identified STIL-associated cancerous mutation, *i.e.*, S76L can impart cancerous phenotypes to dividing cells.

### STIL-BRCA1 axis disrupted in oncogenic STIL-S76L condition

We found that S76L does not affect BRCA1 protein stability, as an expression of RNAi-resistant EGFP-STIL wild type or S76L in cells depleted with endogenous STIL by siRNA was able to rescue BRCA1 protein levels (**Figure 4A**). Accordingly, we observed similar levels of total BRCA1 protein in cells overexpressing S76L, thus mimicking heterozygous oncogenic condition (**Figure 4B**) compared to the STIL wild-type expressing cells. However, careful analysis of BRCA1 localization by fluorescence microscopy revealed a reduction in the number of BRCA1 nuclear foci with a simultaneous increase of centrosome BRCA1 in cells expressing S76L mutant, as compared to wild-type STIL (**Figure 4C-F**). Although the total amount of EGFP-STIL-S76L protein levels observed by immunoprecipitation are comparable to the EGFP-STIL (**Figure S2C**), it does not localize to centrosomes (**Figure S2D,E**), which could be due to the difference in their protein stability as enhanced ubiquitination of S76L was observed (**Figure S2F**).

**Figure 4.**
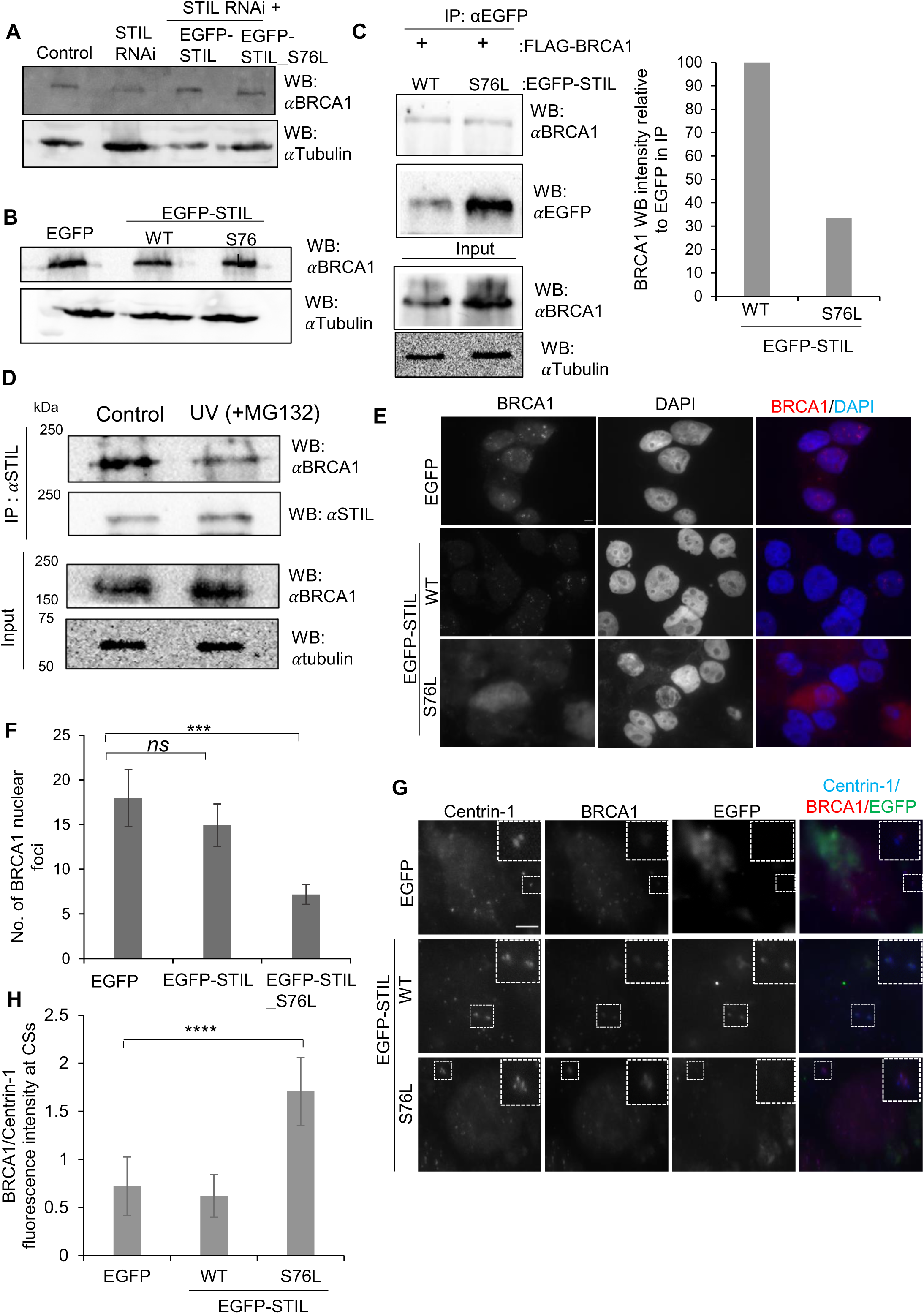
Oncogenic STIL exhibits a disrupted STIL-BRCA1 axis. (**A**) Western blot representing total BRCA protein levels under indicated conditions. α-tubulin serves as a loading control. The blot is representative of two repeats. (**B**) Western blot showing endogenous BRCA1 levels in the total cell lysates of MCF7 cells expressing EGFP (control), EGFP-STIL (WT, wild type; S76L mutant). α-tubulin serves as loading control. The blot is representative of two repeats. (**C**) Immunofluorescence images for S-phase synchronized MCF7 cells immunostained for endogenous BRCA1 and DAPI (nuclear marker) expressing EGFP, EGFP-STIL, and EGFP-STIL-S76L. Scale bar: 5μm. (**D**) Bar graph showing the average number of BRCA1 nuclear foci ±SEM quantified from (C). Results are from two independent experiments (n=70-100). ns, not significant (p>0.05); ***p<0.001 (Two-tailed unpaired Student’s t-test.). (**E**) Immunofluorescence images for S-phase synchronized MCF7 cells immunostained for centrin-1 (centriole marker) and BRCA1. EGFP (green) shows the signal from indicated expressed constructs. Scale bar: 5μm. (**F**) Bar graph representing ratio of BRCA1 *vs*. Centrin-1 fluorescence signal at centrosomes±SEM for respective conditions in (E). Results are from two independent experiments (n=40-50). ****p<0.0001 (Two-tailed unpaired Student’s t-test.)

### BRCA1 is the effector of oncogenic STIL-S76L phenotypes

Since enhanced BRCA1 at S-phase centrosome is linked to DNA damage^47^, we observed more DNA damage in cells overexpressing EGFP-STIL-S76L (**Figure 5A,B**). To restore BRCA1 balance in cells expressing STIL-S76L, we overexpressed FLAG-BRCA1, which was able to rescue the DNA damage phenotype (**Figure 5A,B**), indicating that the loss of BRCA1 nuclear signal as a possible cause of DNA damage in oncogenic STIL. Furthermore, BRCA1 overexpression rescued the amplified centrosome population of oncogenic STIL (**Figure 5C,D**). BRCA1 directly interacts with the centrosome kinase, Aurora-A, and DNA damage inflicted by cisplatin-treatment has been shown to increase both the centrosome BRCA1 and Aurora-A kinase protein levels at S-phase^47^. Accordingly, the expression of S76L oncogenic mutant, which results in DNA damage (**Figure 5A,B**), also has enhanced levels of BRCA1 (**Figure 4E, F**), and Aurora-A levels at S-phase (**Figure 5E,F**). These enhanced Aurora-A levels at centrosome in the oncogenic mutant were rescued on the expression of exogenous FLAG-BRCA1 in the cells. That suggests that BRCA1 is an effector for STIL-dependent regulation of centrosome number and DNA integrity.

**Figure 5:**
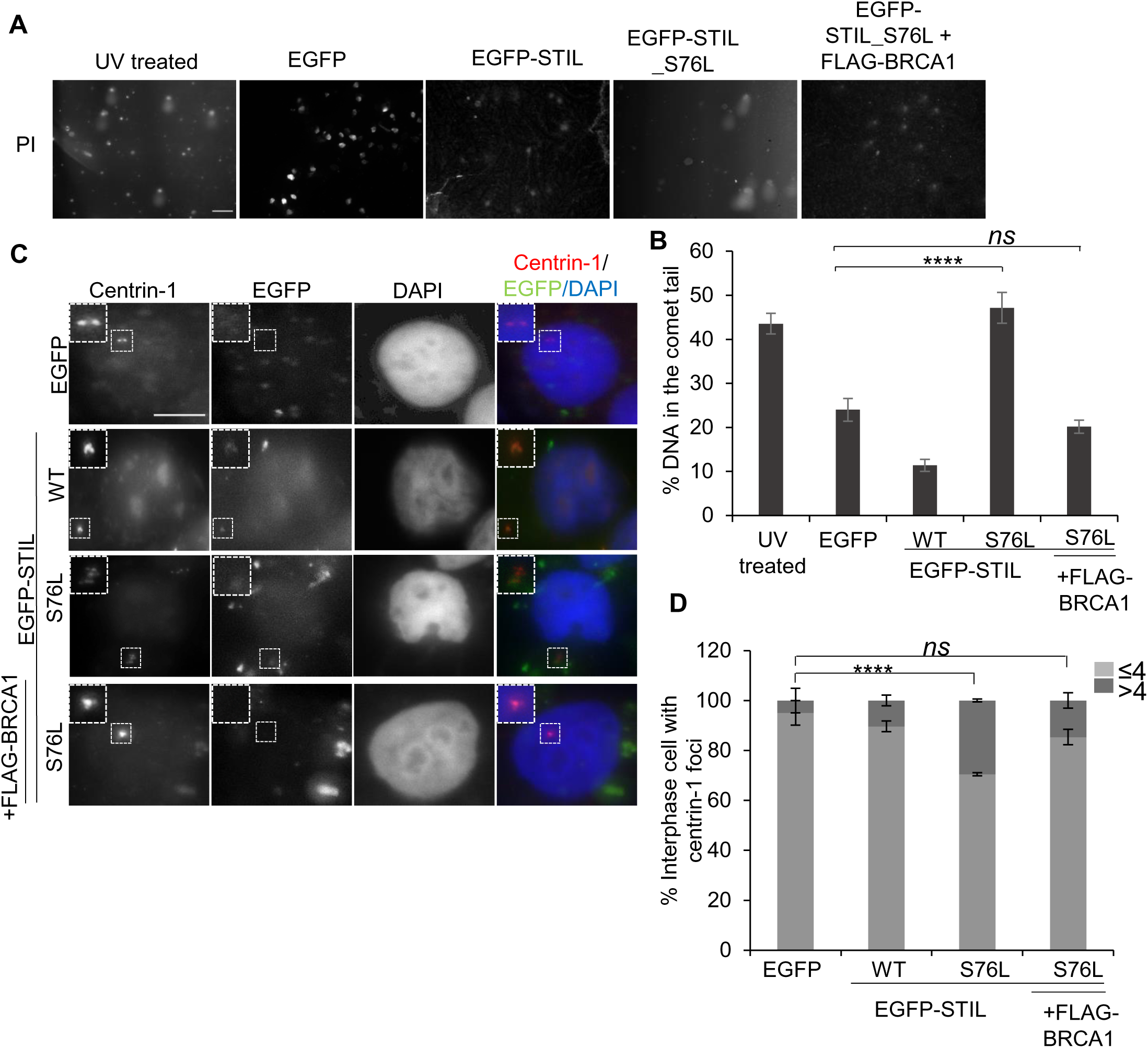
Exogenous BRCA1 restores disrupted STIL-BRCA1 axis. (**A**) Fluorescence images of MCF7 cells expressing EGFP (control), EGFP-STIL, and EGFP-STIL-S67L with DNA stained with propidium iodide (PI) after the comet assay. UV-treated MCF7 cells are a positive control. Scale bar: 100 μm. (**B**) Bar graph showing the average percentage of DNA in the comet tails ±SEM as quantified from (A). Results are from two independent experiments (n=100-375). ns, not significant (p>0.05); ****p<0.0001 (Two-tailed unpaired Student’s t-test). (**C**) Immunofluorescence images of S-phase synchronized MCF7 cells expressing indicated EGFP (control) and EGFP-STIL (WT, wild type; S76L mutant). The last panels show cells co-expressing FLAG-BRCA1. The cells are stained for Centrin-1 (centriole marker) and DAPI (nuclear marker). Insets show a magnified view of the centrioles. Scale bar: 5μm. (**D**) Bar graph showing the mean percentage of interphase cells ±SEM with >4 (dark gray) and ≤4 (light gray) Centrin-1 dots as quantified from (C). Results are from two independent experiments (n=100-150) ns, not significant (p>0.05); ****p<0.0001 (Chi-squared test). (**E**) Immunofluorescence images of S-phase synchronized MCF7 cells expressing indicated EGFP (control) and EGFP-STIL (WT, wild type; S76L mutant). The last panels show cells co-expressing FLAG-BRCA1. The cells are stained for Centrin-1 (centriole marker), Aurora-A (Aur-A), and DAPI (nuclear marker). EGFP signal represents signal from expressed respective constructs viewed in the same channel at centrin-1. Insets show a magnified view of the centrioles. Scale bar: 5μm. (**F**) Bar graph showing the average percentage of DNA in the comet tails ±SEM as quantified from (A). Results are from two independent experiments (n=100-375). ns, not significant (p>0.05); ****p<0.001 (Two-tailed unpaired Student’s t-test).

Despite multiple centrosomes in STIL-S76L expressing cells, multipolar spindles were not apparent. That was due to an increase in the cell population (62.7%) with centrosome clustering by M-phase, which resulted in pseudo-bipolar spindle organization (**Figure 6A,B**). Accordingly, we could show that treating cells with the HSET/KIFC1-specific inhibitor, CW069, enhanced centrosome declustering (**Figure 6C,D**), thus identifying a promising therapeutic target in such oncogenic conditions.

**Figure 6:**
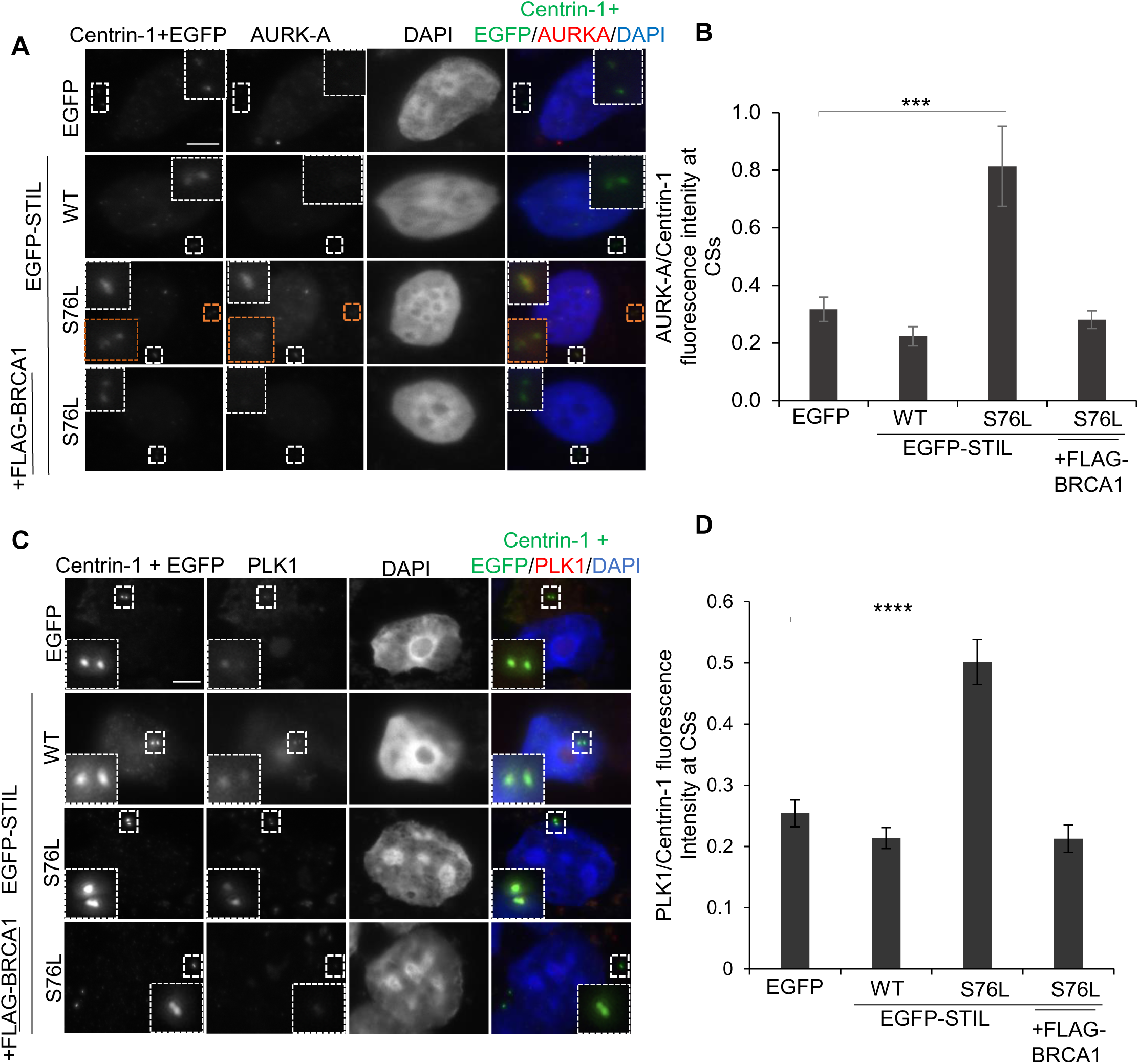
Oncogenic STIL shows kinesin-dependent centrosome clustering. (**A**) Representative images of M-phase synchronized MCF7 cells expressing EGFP (control), EGFP-STIL (WT, wild type; S76L mutant). The cells are immunostained with α-tubulin (red), centrin-1+EGFP signal (green), and DAPI (blue). Scale bar: 5µm. (**B**) The cartoon depicts that cells with more than two Centrin-1 dots per pole were regarded as centriole clustering. Bar graph representing the average percentage (%) of mitotic cells with centriole clustering ±SEM for the indicated condition in (A). (**C**) Representative images of mitotic cells from unsynchronized MCF7 cells expressing EGFP (control), EGFP-STIL, EGFP-STIL_S76L without (-) or with (+) CW069 treatment (50 μM, 6 hrs). The cells are immunostained with Centrin-1 (red) and DAPI (blue). Scale bar: 5µm. Results are from two independent experiments (n=16-22) (**D**) Bar graph representing average percentage (%) of mitotic cells with normal (grey), clustered (red), and declustered (blue) centrioles ±SEM for indicated condition in (C) for the untreated (-) and treated (+) with CW069.

## DISCUSSION

BRCA1 is a well-known tumor suppressor protein involved in DNA damage response pathways, and STIL is a proto-oncogene required for centrosome organization and cell proliferation. STIL can form a complex with BRCA1, but little is understood about their relevance in cell homeostasis. This work establishes the STIL-BRCA1 axis in regulating centrosome number and DNA integrity. We show that double-stranded DNA damages inflicted by either UV treatment or BRCA1 depletion caused STIL-mediated centrosome amplification. Immunoprecipitation shows that the two proteins are part of the same complex and co-localize at centrosomes at the S-phase in breast cancer cells, MCF7. STIL regulates BRCA1 protein stability, as depletion of STIL by siRNA reduced total BRCA1 protein *via* ubiquitination-dependent pathway. Subsequently, we show that BRCA1 depletion causes DNA damage and centrosome amplification phenotypes. We further show that this STIL-BRCA1 axis is disrupted in cancer patients with a heterozygous missense mutation in STIL (S67L). The oncogenic STIL-S76L overexpression enhances centrosome numbers as observed by fluorescence microscopy and cell migration in scratch assay, confirming its oncogenic impact in cells. Although the total BRCA1 levels remain unaffected in oncogenic STIL, the nuclear BRCA1 puncta are reduced with its simultaneous enhanced localization at centrosomes in the S-phase. Accordingly, we observed an increase in DNA damage in these cells. The DNA damage correlates with the enhanced levels of centrosome kinase Aurora-A, which could cause centrosome amplification in oncogenic STIL conditions. The disrupted STIL-BRCA1 axis can be restored by overexpression of exogenous FLAG-BRCA1, as DNA damages and centrosome number phenotypes were rescued. The cells expressing STIL-S76L show centrosome clustering and pseudo bipolar spindles, which is a promising therapeutic target for small molecule inhibitors^48^. We further show that treatment of clustered centrosomes in STIL-S76L with the declustering agent CW069^49^ results in their declustering, which could have promising anti-cancer therapeutic applications. This work reports a novel STIL-BRCA1 axis, which opens new possibilities for chemo-therapeutic cancer interventions.

## MATERIAL & METHODS

### Cell Culture and Synchronization

MCF7 cells were maintained in Dulbecco’s modified Eagle’s medium (Himedia#Al111) supplemented with 1% L-glutamine, 1% Pen/Strep, and 10% fetal bovine serum (Himedia #RM10434,). Cells were synchronized in the S phase using a double-thymidine block^50^. Briefly, cells were incubated with 2 mM thymidine (Sigma# T9250) for 18 hrs, followed by 9 hrs of release, and the second block was introduced for 15 hrs, followed by harvesting. For the M-phase, cells were synchronized using a double-thymidine block followed by a 12 hrs release.

### siRNA, Plasmids and Transfection

Double-stranded STIL siRNA and BRCA1 siRNA oligonucleotides were synthesized (Biotech Desk Pvt Ltd) with 3’-UU overhangs with the following sense strand sequences, 5’-GUUUAAGGGAAAAGUUAUU-3’^51^ and 5’-CAGGAAAUGGCUGAACUAGAA-3’^52^, respectively. siRNA (72 nM) was transfected using Lipofectamine 3000 reagent as per the manufacturer’s protocol (Invitrogen #L3000001). The pcDNA5 FRT/TO MycLAP-STIL plasmids was ordered from Addgene (#80266). pcDNA5 FRT/TO MycLAP-STIL_S76L and pcDNA5 FRT/TO MycLAP-STIL_S76L_S183A were generated using site-directed mutagenesis, using primers 5’-CAGAATAAAAAAAATTTGTCATGCTTTTTACTT-3’ and 5’-GAGAGTTTGGACGCTGTGGAATTT-3’ respectively. pDEST 3xFlag-pcDNA5-FRT/TO-BRCA1 was ordered from Addgene (#52504) Plasmids were transfected using Polyethylenimine Max^53^ (Polysciences #23966-100).

### Primary & Secondary antibodies

Following primary antibodies for immunofluorescence (IF) and western blotting (WB) with indicated dilution were used: mouse anti-γ tubulin antibody (Sigma #T5326, 1:100 IF), rabbit anti-Centrin-1 antibody (Abcam#ab101332, 1:500 IF), mouse anti-Centrin-1 20H5 antibody (Merck Millipore#04-1624, 1:200 IF), mouse anti-α tubulin antibody (Sigma#T9026, 1:100 IF, 1:1000 WB), mouse anti-BRCA1 D-9 antibody (Santa Cruz Biotechnology#SC6954, 1:100 IF, 1:1000 WB), rabbit anti-STIL antibody (Abcam#ab89314, 1:50 IF, 1:2000 WB), rabbit anti-ubiquitin antibody (Abcam#ab134953, 1:2000 WB), mouse anti-phosphorylated p53 (CST#9286T, 1:1000 WB), mouse anti-p53 (CST#2524S, 1:1000 WB), mouse anti-p21 (CST#2946T, 1:1000 WB), and rabbit anti-GFP antibody (Sigma #G1544, 1:7000 WB). Following secondary antibodies at the indicated dilution were used for IF: Anti-rabbit IgG (Fc) TRITC (Sigma #SAB3700846, 1:500), Anti-mouse IgG (Fc) TRITC (Sigma #SAB3701020, 1:500), Anti-rabbit IgG FITC (Sigma #F7512, 1:750), Anti-mouse IgG FITC (Sigma #F5387, 1:750). Following secondary antibodies at the indicated dilution were used for WB: Anti-mouse IgG HRP antibody (Cell Signaling Technology #7076S, 1:10000) and anti-rabbit IgG HRP (Cell Signaling Technology #7074S, 1:10000).

### Immunofluorescence Microscopy

Cells were seeded on a poly-D-lysine hydrobromide (P7280, Sigma) coated 12 mm coverslips. They were permeabilized for 2 minutes using 0.1% Triton X in 1X PBS and fixed using 4% paraformaldehyde for 10 minutes. After three washes with 1X PBS, blocking solution (1% BSA in PBS + 0.1% Tween 20) was added for 1 hr. Primary antibodies were diluted in a blocking solution, and the coverslips were kept at 4°C overnight. This was followed by three washes with 1X PBS and incubation with secondary antibody solutions for 1 hr at room temperature. After three washes with 1X PBS, 0.5μg/ml DAPI in 1X PBS was added to the cells for 10 minutes in the dark to stain the nucleus. The coverslips were mounted onto a glass slide using DABCO-Mowiol mounting media and stored at 4°C. Fluorescence images were taken using an Olympus IX83 fluorescence microscope with 100X oil, 1.35 NA objective lens. The images were analyzed using the ImageJ software.

### Immunoprecipitation and Western blotting

Cells were suspended in ice-cold RIPA buffer (150mM NaCl, 5mM EDTA pH 8, 50mM Tris pH 8, 1% NP40, 0.5% sodium deoxycholate, 0.1% SDS) supplemented with 0.5% protease and phosphatase inhibitors and lysed by passing through a small syringe needle. Lysates were cleared by centrifugation at 14,500xg for 30 minutes. The cell lysate was incubated with an anti-STIL antibody (1:50) or anti-GFP antibody (1:200) overnight at 4°C. Protein G-sepharose beads were added to the lysate-antibody mixture for 4 hours at 4°C and kept for shaking. Beads were washed with 1X RIPA thrice at 300xg for 2 minutes. The protein complex was eluted in the Laemmli sample buffer and resolved on 10% polyacrylamide gels. For western blotting, resolved protein bands from SDS-PAGE were transferred to the nitrocellulose membrane. The membrane was blocked using 5% BSA in 1xPBST (1X PBS with 0.05% Tween-20) for 1 hr and incubated with primary antibody at 4°C overnight. The membrane was washed with 1X PBST thrice and incubated with the HRP-conjugated secondary antibody for 1 hour at room temperature. The blot was developed using Clarity Western ECL substrate (Biorad #1705061) for 5 minutes in the dark and analyzed using gel documentation system (Biotron Omega Lum G).

### Cell Migration Assay

MCF7 cells at 50% confluency were transfected with EGFP-STIL constructs. At 70% confluency, the cells were given a media change to serum-free DMEM. The monolayer was scratched with a pipette tip at nine different reference points, and images were taken at 0 hr, 24 hrs, and 48 hrs time points using the Olympus IX83 fluorescence microscope with a 20X dry objective^54^. Images were analyzed using the Wound Healing Size ImageJ plugin^55^. The percentage of wound closure was calculated as follows –

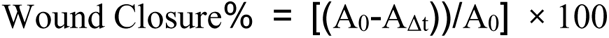

Where A_0_ is the initial wound area, A_Δt_ is the wound area after n hours of the initial scratch.

### Ubiquitination Assay

MCF7 cells were transfected with STIL constructs or respective RNAi and synchronized in S-phase using a double thymidine block. 5μM MG-132 was added to the cells for 16 hours before lysis. Cells were lysed in RIPA buffer supplemented with 0.5% protease and phosphatase inhibitors and 10mM N-ethylmaleimide. Immunoprecipitation was performed on the respective primary antibody. Protein complexes were blotted on a nitrocellulose membrane and incubated with rabbit anti-ubiquitin primary antibody (Abcam # ab134953) to look at protein ubiquitination in a western blot.

### Comet Assay

Agarose-coated slides were prepared by dipping in 1% agarose solution in 1X PBS. In the case of UV exposure condition, the MCF7 cells were exposed to UVC for 30 minutes. For endogenous DNA damage, MCF7 cells were transfected with the indicated plasmid constructs for 48 hrs, and 2 X 10^5^ cells/ml were added to a 24-well plate. 1 ml of 1% Low Melting Point agarose solution in 1X PBS was added to each well and mixed with cells gently. The mixture was overlaid on agarose-coated slides on ice and kept for 15 minutes. All steps after this were performed in the dark. The slides were incubated in the lysis buffer (2.5M NaCl, 100mM EDTA, 10mM Tris, 200mM NaOH at pH 10.5, 1% Triton-X100, and 1% DMSO) on ice for 2 hours. Subsequently, the slides were incubated in cold electrophoresis buffer (300mM NaOH, 1mM EDTA, 1% DMSO at pH > 13) for 30 minutes. Slides were shifted to an electrophoresis chamber and run at 21V for 30 minutes. After two washes with distilled water, the slides were stained using 2.5 mg/ml Propidium iodide solution. The excess of stain was washed with distilled water, and the slides were dried overnight at room temperature. The images were captured using a 10X objective of the Olympus IX83 fluorescence microscope. DNA damage was assessed using the OpenComet plugin available for ImageJ^56^.

### Centriole counting and Fluorescence Intensity Quantification

Centrin-1 was used as a centriolar marker for the centriole counting. The number of centrin-1 dots were counted for each cell and the number of cells with ≤ 4 or >4 centrioles were scored. The percentage of cells with ≤ 4 or >4 centrioles was calculated by dividing by the total number of cells. The mean percentage±SEM was calculated from 2-3 independent experiments.

For intensity measurements, the Centrin-1 signal was used as a reference to draw a region of interest (ROI) in ImageJ^57^. The same ROI was duplicated into the EGFP channel. The exact size ROI was also placed around the centrosome to measure the background. The ratio of absolute fluorescence intensity of EGFP *vs*. Centrin-1 was reported after subtracting the average background signal of the respective channel.

### Centrosome declustering assay

MCF7 cells were seeded on poly-D lysine-coated coverslips and transfected with respective EGFP constructs for 48 hours. 6 hrs before analysis 50μM CW069 (HSET-selective inhibitor) was added to treated cells. Cells were immunostained using Centrin-1 as a centriolar marker. Mitotic cells with background EGFP signals were scored. Cells with 2 centrosomes (with 2 centrin-1 dots per centrosomes) were considered normal phenotypes. Cells with more than 2 centrin-1 dots per centrosome were considered as clustered phenotype. Cells with more than 4 separated centrioles centrin-1 dots were considered declustered.

### Statistical analysis and Software

The bar graphs were generated in Microsoft Excel. Experiments were repeated at least twice. For intensity measurements, the p-value was calculated using a two-tailed Student’s t-test with unequal variance for the two conditions as indicated on top of bar graphs using a solid line. The p-values for centriole and spindle pole counting experiments were calculated using the chi-square test. Figure panels were arranged on Microsoft PowerPoint.

## SUPPLEMENTAL INFORMATION

Figure S1 and Figure S2

## DATA AVAILABILITY

This study includes no data deposited in external repositories

## ACKNOWLEDGEMENTS

The pcDNA5 FRT/TO/EGFP plasmid is a gift from the laboratory of Prof. Andrea Mussachio, MPI Dortmund, Germany. We thank the Centre for Research & Development of Scientific Instruments at IIT Jodhpur for the microscopy facility.

## FUNDING

The work is supported by the grants received from the Board of Research in Nuclear Sciences (55/14/02/2021-BRNS/10206) and the Department of Biotechnology (BT/12/IYBA/2019/02)

## DISCLOSURE & COMPETING INTERESTS STATEMENT

The authors declare no competing or financial interests.

## AUTHOR CONTRIBUTIONS

Conceptualization: S.S., P.S.; Methodology: S.S.; Validation: S.S., P.S.; Formal analysis: S.S., P.S.; Investigation: S.S.; Data curation: S.S., P.S.; Writing original draft: S.S. P.S.; Writing-review & editing: P.S.; Supervision: P.S.; Project administration: P.S.; Funding acquisition: P.S.

---

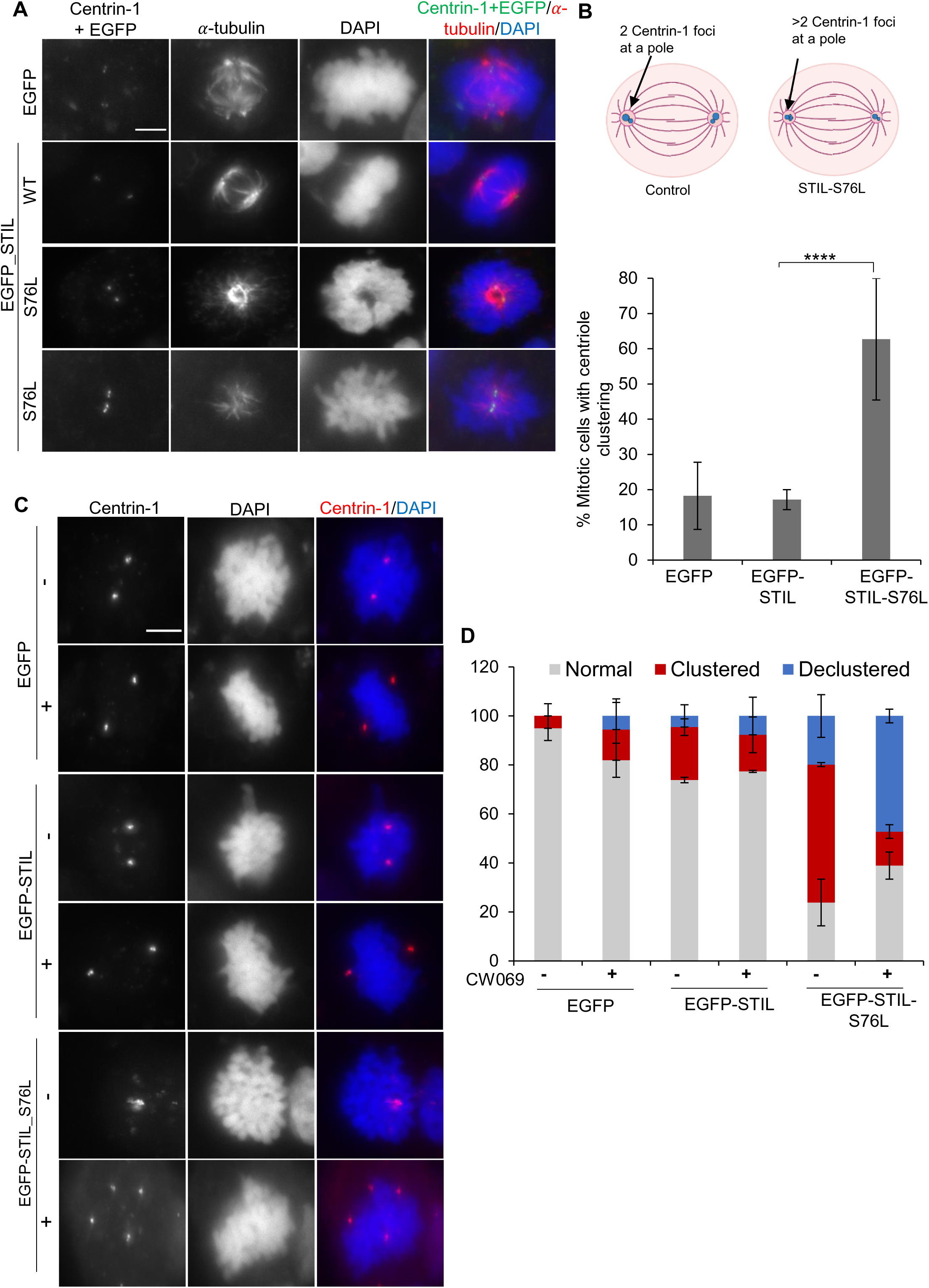

## Notes

### Competing Interest Statement

The authors have declared no competing interest.

### Summary of Updates

The version of this manuscript has been revised to include additional experiments and analysis which has helped us to refine the proposed model.

