## Supplementary material for "Oncogenic STIL-mediated loss of BRCA1 functionality causes DNA damage and centrosome amplification": S1, S2 and S3

Author Affiliations:

*Running title*: STIL-BRCA1 axis in cancer

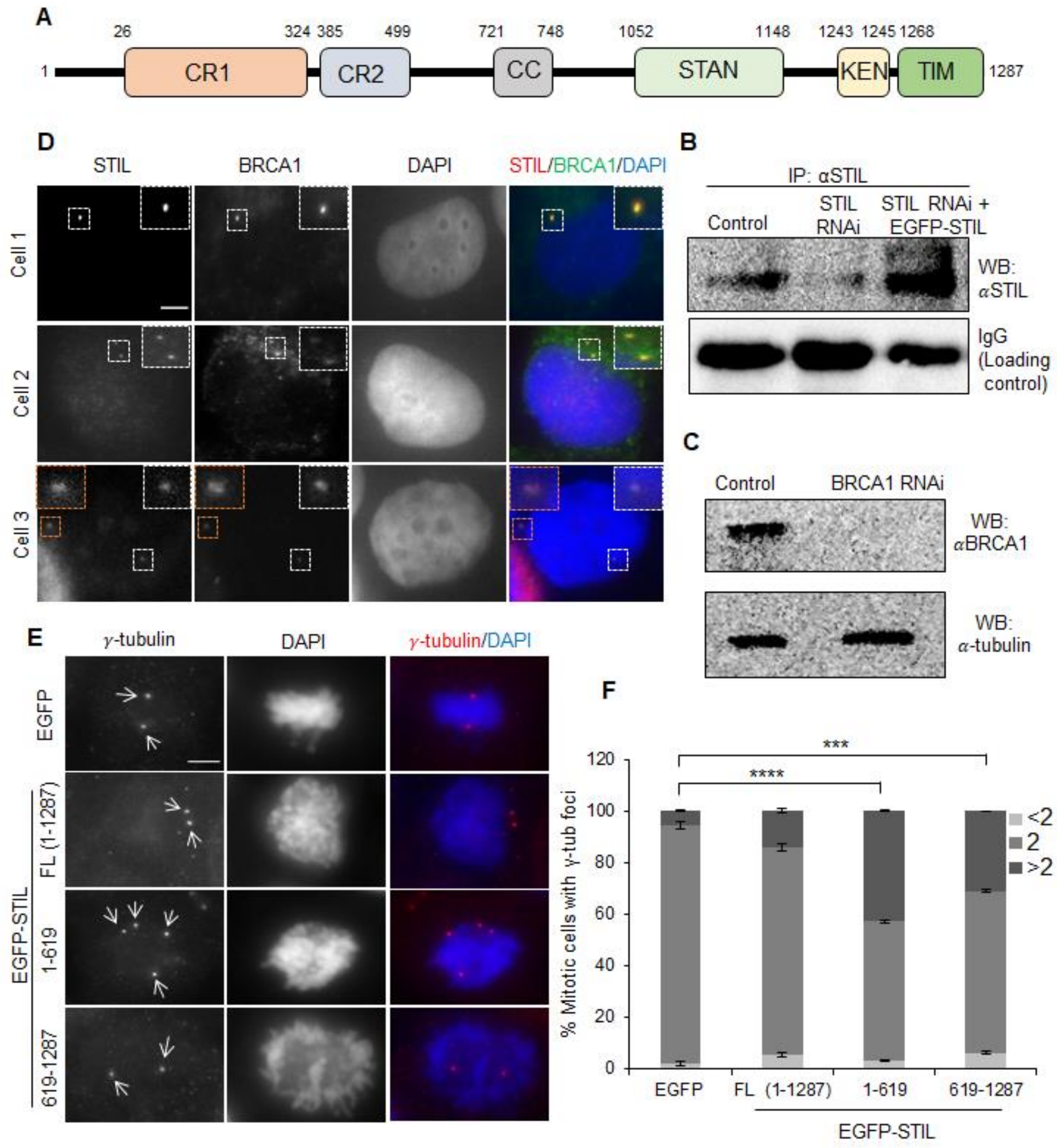


**Figure S1**: (**A**) Schematic representation of conserved regions of human STIL. Numbers indicate amino acid residues marking the position of different regions in the protein. CR1 (conserved region 1), CR2 (conserved region 2), and CC (coiled-coil). (**B**) Immunofluorescence images of S-phase synchronized MCF7 cells stained for endogenous STIL & BRCA1. DAPI stains the nucleus. Insets show a magnified view of centrioles. Scale bar: 5μm. (**C**) Western blot showing immunoprecipitated (IP) STIL levels in control, STIL RNAi (72 nM, endogenous STIL depletion), and STIL RNAi+RNAi-resistant EGFP-STIL condition. IgG serves as a loading control. (**D**) Western blot showing BRCA1 depletion after RNAi-treatment (72 nM) in the total cell lysate of MCF7 cells. α-tubulin serves as loading control. (**E**) Immunofluorescence images of mitotic MCF7 cells transfected with EGFP (control) or EGFP-STIL [full-length, FL (1-1288 residues); 1-619 residues (N-terminal); 619-1288 (C-terminal)] and stained with pericentriolar marker, γ-tubulin and nuclear stain, DAPI. White arrows mark centrosomes/cells. Scale bar: 5μm. (**F**) Bar graph showing the mean percentage of mitotic cells±SEM with >2 (dark gray), 2 (medium gray), and <2 (light gray) γ-tubulin dots in the respective conditions. Results are from two independent experiments (n=50-100). ***$p<0.001$ and ****$p<0.0001$ (Chi-squared test).

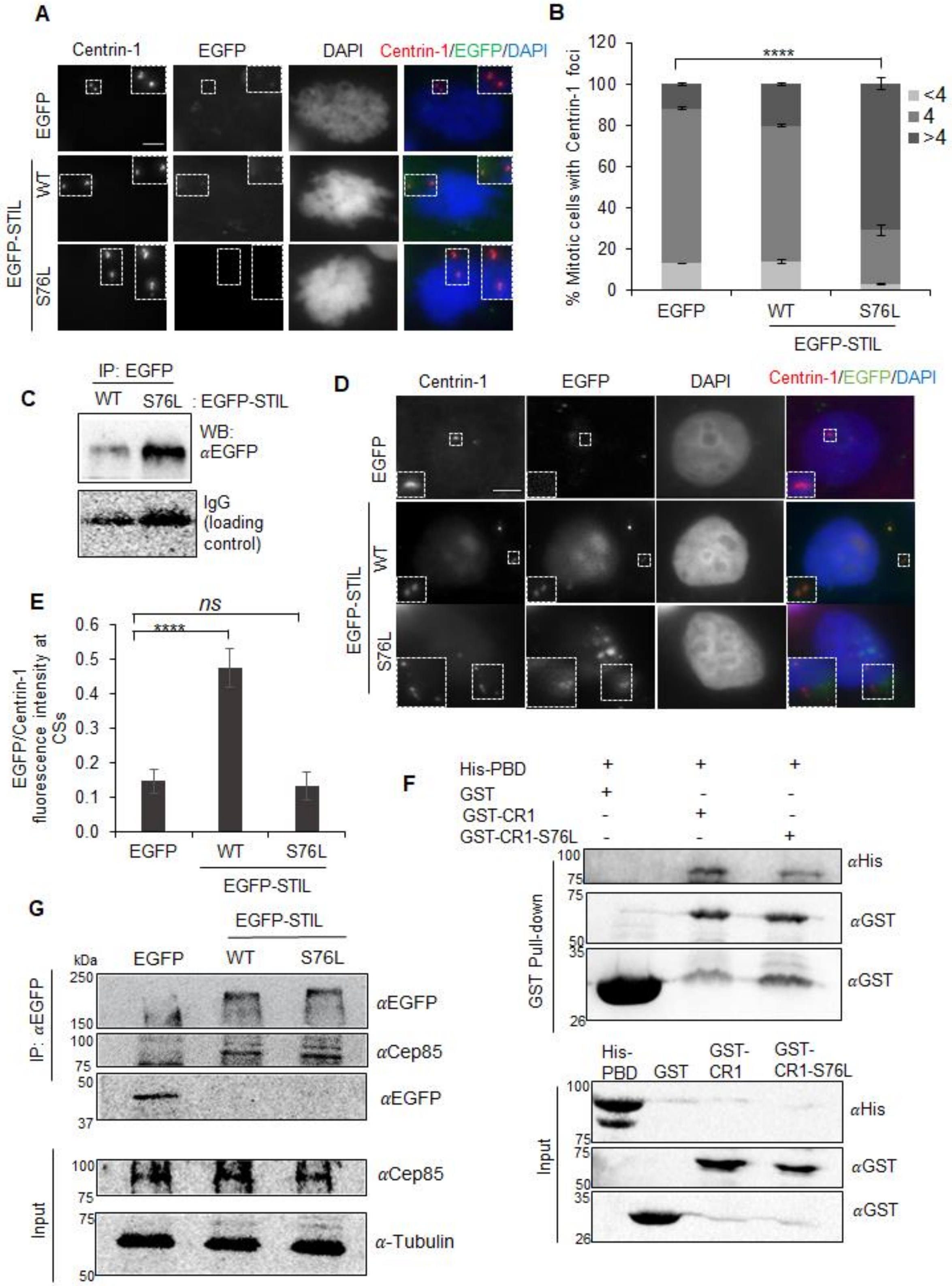
A
Centrin-1
EGFP
DAPI
Centrin-1/EGFP/DAPI
EGFP
WT
S76L
EGFP-STIL
B
% Mitotic cells with Centrin-1 foci
****
<4
4
>4
EGFP
WT
S76L
EGFP-STIL
C
IP: EGFP
WT
S76L
: EGFP-STIL
WB: αEGFP
IgG (loading control)
D
Centrin-1
EGFP
DAPI
Centrin-1/EGFP/DAPI
EGFP
WT
S76L
EGFP-STIL
E
ns
****
EGFP/Centrin-1 fluorescence intensity at CSs
EGFP
WT
S76L
EGFP-STIL
F
His-PBD
GST
GST-CR1
GST-CR1-S76L
αHis
αGST
αGST
GST Pull-down
His-PBD
GST
GST-CR1
GST-CR1-S76L
Input
G
EGFP-STIL
kDa
EGFP
WT
S76L
IP: αEGFP
αEGFP
αCep85
αEGFP
Input
αCep85
α-Tubulin

**Figure S2:** (**A**) Immunofluorescence images of mitotic MCF7 cells expressing EGFP (control) or EGFP-STIL (wild type, WT; S76L, mutant). The cells are stained with centrin-1 (centriolar marker) and DAPI (nuclear marker). EGFP signal represented for respective constructs. Insets show a magnified view of centrioles. Scale bar: 5μm. (**B**) Bar graph showing the mean percentage of mitotic cells±SEM with >4 (dark gray), 4 (medium gray), and <4 (light gray) centrin-1 dots in the respective conditions in (F). Results are average from three independent experiments (n=50-100). ****p<0.0001 (Chi-squared test). (**C**) Western blot showing immunoprecipitated EGFP-STIL and EGFP-STIL from the total cell lysate of S-phase synchronized MCF7 cells using EGFP antibody. IgG bands serve as loading controls. The blot is representative of two repeats. (**D**) Immunofluorescence images of S-phase synchronized MCF7 cells expressing EGFP (control) or EGFP-STIL (wild type, WT; S76L, mutant). The cells are stained with centrin-1 (centriolar marker) and DAPI (nuclear marker). EGFP signal represented for respective constructs. Insets show a magnified view of centrioles. Scale bar: 5μm. (**E**) Bar graph showing the ratio of EGFP *vs*. centrin-1 fluorescence signal±SEM from the experiment (D). Results are from three independent experiments (n=74-94). ns, not significant (p>0.05); ****p<0.0001 (Two-tailed unpaired Student's t-test). (**F**) Western blot showing interaction between the GST alone (tag control), GST tagged CR1 region of STIL (wild type and S76L mutant), and His tagged Polo-box domain (PBD) of PLK4 using the GST pull-down assay. (**G**) The western blot shows the interaction of EGFP-STIL (wild type and S76L mutant) with CEP85 by immunoprecipitation assay using an antibody against EGFP. The lower panel shows CEP85 input, and α-tubulin serves as loading control.

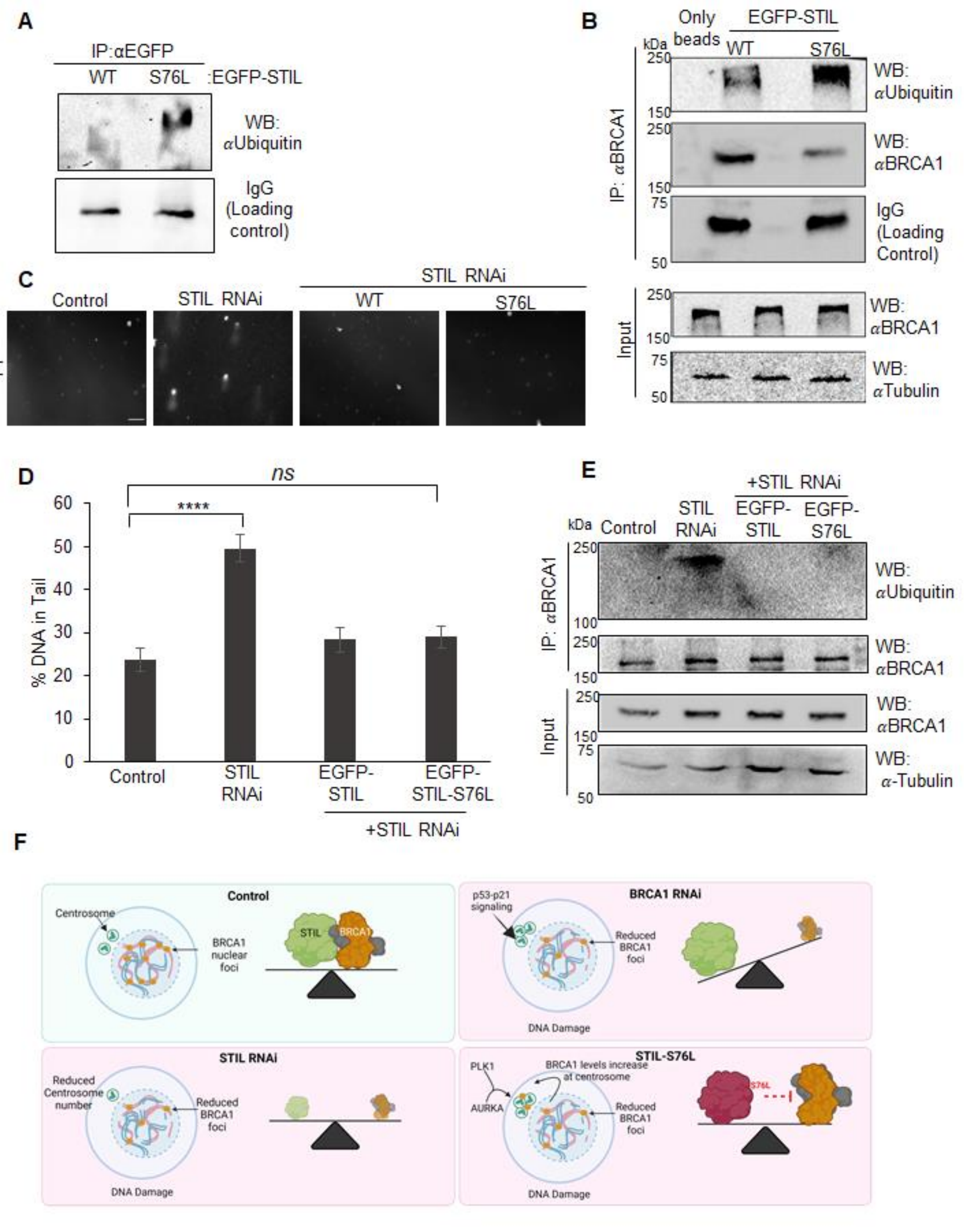


**Figure S3**: (**A**) Western blot representing immunoprecipitated (IP) EGFP-STIL and EGFP-STIL-S76L using the EGFP antibody. The blot is developed using an antibody against ubiquitin.

IgG serves as loading control. The blot is a representation of two repeats. (**B**) The western blot shows ubiquitination of BRCA1 after IP from cells expressing EGFP-STIL (wild type and S76L mutant) using an antibody against BRCA1. The blot was developed using antibodies against Ubiquitin and BRCA1. There is no non-specific binding to lysates to only beads. IgG serves as loading control for IP. The lower panels represent BRCA1 input, and α-tubulin serves as a loading control. (**C**) The comet assay shows DNA damage in control, STIL depleted (RNAi), and STIL RNAi complemented with either EGFP-STIL wild type (WT) or S76L. Scale bar: 100 μm. (**D**) Bar graph showing the average percentage of DNA in the comet tails ±SEM as quantified from (C). Results are from two independent experiments (n=150-200). ns, not significant ($p>0.05$); ****$p<0.0001$ (Two-tailed unpaired Student's t-test). (**E**) The western blot showing ubiquitination of BRCA1 after IP from control, STIL depleted (RNAi), and STIL RNAi complemented with either EGFP-STIL wild type or S76L cells using an antibody against BRCA1. The blot was developed using antibodies against Ubiquitin and BRCA1. The lower panels represent BRCA1 input, and α-tubulin serves as a loading control. (**F**) The proposed model depicts that STIL interacts with the BRCA1 protein complex and regulates DNA integrity by maintaining nuclear signals of BRCA1, which in turn is required for regulating centrosome number. STIL depletion results in DNA damage as the BRCA1 protein stability is affected.

However, the centrosome numbers are reduced, as STIL is a core centrosome protein. The BRCA1 depletion elicits p53-p21-dependent centrosome amplification. In the case of oncogenic STIL-S76L, interaction with BRCA1 is disrupted due to the DNA damage signaling, which subsequently translocates BRCA1 to centrosomes. This enhances centrosome levels of Aurora-A and BRCA1, causing centrosome amplification.
